# TPX2 activation by GM130 controls astral microtubule formation and spindle orientation

**DOI:** 10.1101/2020.03.16.994426

**Authors:** Haijing Guo, Jen-Hsuan Wei, Joachim Seemann

## Abstract

Spindle orientation is important in multiple developmental processes as it determines cell fate and function. The correct orientation of the mitotic spindle depends on the assembly of a proper astral microtubule network. Here, we report that the spindle assembly factor TPX2 regulates the astral microtubule network. TPX2 in the spindle pole area is activated by GM130 on mitotic Golgi membranes to promote astral microtubule growth. GM130 relieves TPX2 inhibition by competing for importin α binding. During mitosis, phosphorylation of importin α at Serine 62 by Cdk1 switches its substrate preference from TPX2 to GM130, thereby enabling competition-based activation. Importin α S62A impedes local TPX2 activation and compromises astral microtubule formation, ultimately resulting in misoriented spindles. Blocking the GM130-importin α-TPX2 pathway impairs astral microtubule growth. Our results reveal a novel role for TPX2 in the organization of astral microtubules. Furthermore, we show that the substrate preference of the important mitotic modulator importin α is regulated by Cdk1 phosphorylation.

## Introduction

Cell proliferation requires accurate partitioning of all vital cellular constituents during cell division. In eukaryotic cells, the microtubule-based mitotic spindle machinery delivers a precise set of chromosomes into each daughter cell. Spindle microtubule filaments are further responsible for the correct positioning and orientation of the spindle to define the cell division plane, which plays a critical role in cell fate decision, morphogenesis and maintenance of tissue organization (Bergstralh et al., 2017). Consequently, aberrant spindle orientation causes defects in various physiological processes such as gastrulation, neuronal differentiation, epithelial self-renewal and tissue stratification (Gong et al., 2004; Lechler and Fuchs, 2005; Fish et al., 2006; Cabernard and Doe, 2009). Determination and maintenance of spindle orientation largely rely on the mitotic microtubule network and its connections to the cell cortex.

Assembly of the mitotic spindle is tightly controlled in space and time through regulated microtubule nucleation and polymerization (Prosser and Pelletier, 2017). In higher eukaryotes, mitotic microtubule arrays predominantly originate at the centrosomes, which form the spindle poles. Astral microtubules emanate from the spindle poles and extend to the cell cortex. This spindle-cortex connection is required to generate the pulling forces that dictate positioning and orientation of the spindle. Reductions of astral microtubules weaken connections between the spindle and the cell cortex, which can often lead to spindle oscillation and misorientation (Petry, 2016). Though astral microtubules are important in spindle positioning, microtubule filaments connected to kinetochores are directly engaged in the segregation of chromosomes into the daughter cells. Kinetochore microtubules are bundles of parallel microtubule filaments known as K-fibers that originate at the spindle poles and attach to the chromosomes via kinetochores. Proper attachment of all K-fibers in the bipolar spindle is required to silence the spindle assembly checkpoint and allow progress into anaphase (Prosser and Pelletier, 2017).

In early mitosis, kinetochore microtubules are nucleated at the chromatin. This key step of spindle formation ensures that all chromosomes are captured by the kinetochore microtubules, with this process being regulated by the small GTPase Ran. Upon loading with GTP (via its chromatin-localized guanine nucleotide exchange factor RCC1), RanGTP destabilizes ternary complexes of classical nuclear localization sequence (NLS)-containing proteins plus importin α/ß and promotes their dissociation (Carazo-Salas et al., 1999). Through this mechanism, RanGTP relieves importin α-mediated inhibition of essential spindle assembly factors to promote microtubule nucleation, stabilization and spindle organization (Kalab and Heald, 2008). One of the key spindle assembly factors is the microtubule nucleator TPX2. Once released from importin α inhibition, TPX2 activates and stabilizes active Aurora A kinase by shielding it from dephosphorylation (Bayliss et al., 2003). The Aurora A-TPX2 complex binds to microtubules, further enhancing microtubule polymerization at the chromosomes (Kufer et al., 2002; Anderson et al., 2007).

In addition to functioning at chromatin, the microtubule nucleation activity of TPX2 can also occur at mitotic Golgi membranes through the Golgi-resident membrane protein GM130 (Wei et al., 2015). During mitosis, the single copy of the mammalian Golgi apparatus is disassembled into a collection of vesicles that cluster and localize in the spindle pole area (Shima et al., 1998). These mitotic Golgi clusters are subsequently delivered by the spindle into the daughter cells where the membranes fuse to re-establish a contiguous Golgi structure (Wei and Seemann, 2017; Mascanzoni et al., 2019). Mitotic disassembly of the Golgi is initiated by Cdk1-mediated phosphorylation of the Golgi-resident membrane protein GM130 (Lowe et al., 2000), which exposes its NLS motif that sequesters importin α from TPX2 by direct competition. TPX2 is thereby activated to locally drive microtubule nucleation at mitotic Golgi membranes. The nascent microtubules are then captured by GM130, which couples Golgi membranes to the spindle (Wei et al., 2015). However, it had remained unclear how TPX2 is preferentially liberated from importin α by the NLS motif of GM130, given the high abundance of NLS-containing proteins that are released upon nuclear envelope breakdown during mitotic entry. Furthermore, how importin α regulated this event spatiotemporally so that the process could be coordinated with other microtubule nucleation activities to form a functional spindle had remained elusive. Moreover, the exact role of Golgi-based microtubule nucleation in shaping the mitotic microtubule network had not been characterized.

Here, we describe a pathway responsible for driving astral microtubule formation and spindle orientation. We show that TPX2 promotes astral microtubule growth around the spindle poles through Aurora A kinase. TPX2 becomes locally activated by GM130, which sequesters importin α from TPX2 on mitotic Golgi membranes around the spindle poles. This sequestration-activation process is triggered by Cdk1-mediated phosphorylation of importin α at Ser62. Mitotic phosphorylation of importin α weakens its interaction with TPX2 but enhances its binding to GM130, thereby enabling competition-based activation of TPX2. This Golgi-based TPX2 activation is required for astral microtubule growth from the spindle poles, since suppressing importin α phosphorylation results in diminished astral microtubule filaments and misorientation of the spindle.

## Results

### Importin α is mitotically phosphorylated by Cdk1 at serine 62

During mitosis, importin α silences several essential spindle assembly factors (such as TPX2) in the cytoplasm by binding to their nuclear localization sequence (NLS). It had been unclear what mechanism enables importin α to preferentially recognize these spindle assembly factors in the presence of other abundant NLS-containing proteins that are released upon nuclear breakdown at mitotic entry. Previous studies showed that post-translational modifications of importin α, such as phosphorylation by casein kinase 2, can modulate its activity (Hachet et al., 2004; Brownlee and Heald, 2019). We hypothesized that mitotic phosphorylation enables importin α to specifically regulate spindle assembly factors during M-phase. To examine if importin α is mitotically modified, we separated lysates from interphase and mitotic HeLa cells on a Phos-tag acrylamide gel, on which phosphorylated proteins migrate more slowly (Figure 1A). By immunoblotting, we observed that the band corresponding to importin α from mitotic lysate was shifted upward relative to that of interphase lysate, as was the positive control GRASP55 (Xiang and Wang, 2010). To further establish if the decreased mobility of mitotic importin α was induced by phosphorylation, we incubated mitotic lysate with λ phosphatase before analyzing the gel-shift response. Phosphatase treatment abolished the band shift of both GRASP55 and importin α, demonstrating that importin α is indeed mitotically phosphorylated. Next, we applied mass spectrometry to identify mitosis-specific phosphorylation sites of importin α. To this end, we generated a HeLa cell line that stably expresses FLAG-tagged importin α in a doxycycline-inducible manner, and then isolated tagged importin α from the interphase and mitotic cell lysates. Mass spectrometry analysis revealed a unique phosphorylated site for mitotic importin α, which is located on a fragment comprised of residues 52-68 (Figure 1B).

**Figure 1.**
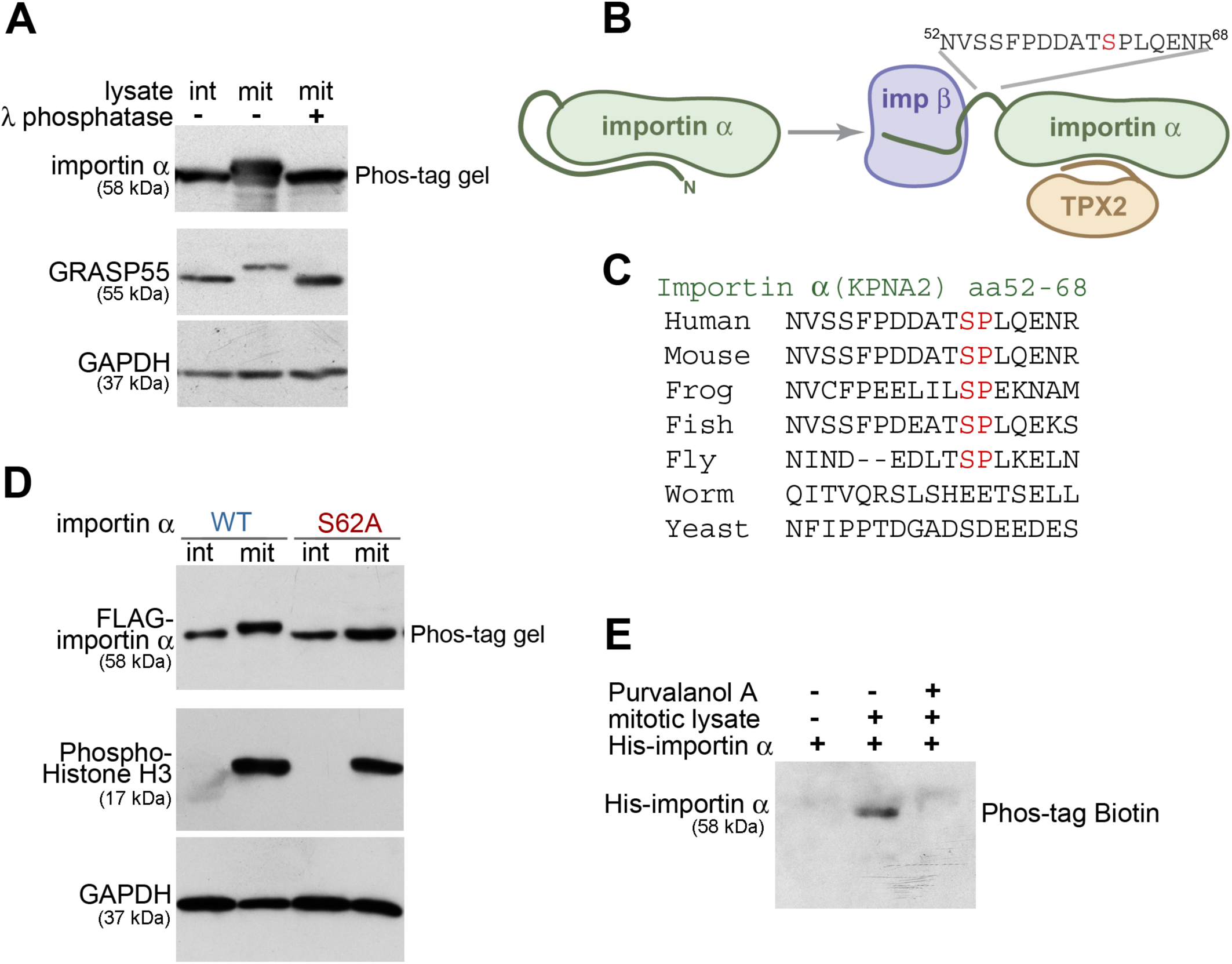
Importin α is mitotically phosphorylated at serine 62 by Cdk1. (A) Importin α is phosphorylated during mitosis. Cell lysates from interphase (int) and mitotic (mit) HeLa cells, as well as λ phosphatase-treated mitotic lysate, were separated on a 10% SDS gel containing 25 μM Phos-tag acrylamide and immunoblotted for importin α. GRASP55 and GADPH were separated by regular SDS-PAGE. (B) Schematic illustration of the location of the mitotically-phosphorylated peptide identified by mass spectrometry. The identified peptide (residues 52-68) harbors the mitosis-specific phosphorylation site of importin α and is located between the importin β-binding domain and the NLS-binding domain. (C) Within the identified peptide sequence, residue serine 62 is conserved among higher eukaryotes. The serine/proline sequence resembles a Cdk1 phosphorylation consensus motif. (D) Importin α is phosphorylated at serine 62. Interphase or mitotic cell lysates from cells expressing FLAG-tagged WT importin α or importin α S62A were separated on a Phos-tag acrylamide gel and immunoblotted for FLAG. Phospho-Histone H3 and GAPDH were separated by SDS-PAGE. (E) Importin α phosphorylation is blocked by the Cdk1 inhibitor Purvalanol A. Beads coated with recombinant full-length His-tagged importin α were incubated with mitotic HeLa extract in the presence or absence of Purvalanol A. The beads were then washed by high salt buffer (PBS, 1M NaCl), and importin α was eluted by boiling in SDS sample buffer. Phosphorylation was detected by blotting with Phos-tag biotin and streptavidin-HRP.

Importin α binds its substrates and forms a ternary complex with importin ß. In the absence of NLS-containing substrates, the autoinhibitory domain (IBB) of importin α folds back into its NLS-binding groove (Figure 1B). Importin ß can bind to the IBB domain of importin α to open up its NLS-binding region. The phosphorylated fragment we identified (residues 52-68) is situated between the IBB domain and the substrate-binding domain. This importin α peptide (NVSSFPDDATSPLQENR) contains several Ser and Thr residues as potential phosphorylation sites, but harbors only one Ser/Thr-Pro consensus motif for the master mitotic kinase Cdk1-cyclinB1 (at Ser62). Since this Ser-Pro motif is highly conserved (Figure 1C), we reasoned that Ser62 is mitotically phosphorylated by Cdk1. To validate this supposition, we generated stable HeLa cell lines that inducibly express FLAG-tagged wild type importin α (WT) or the importin α S62A mutant protein, and then analyzed their interphase and mitotic lysates on a Phos-tag gel (Figure 1D). We found that the mobility of WT importin α from mitotic lysate was reduced compared to that of interphase lysate, whereas the S62A mutant from mitotic lysate migrated at the same speed as from interphase lysate. Next, we determined if importin α is mitotically phosphorylated by Cdk1. We incubated recombinant importin α with mitotic extract in the presence or absence of the Cdk1 inhibitor Purvalanol A. Phosphorylation was then detected by blotting with biotinylated Phos-tag combined with streptavidin-conjugated HRP (Figure 1E). Phos-tag biotin specifically labeled the phosphorylated importin α in mitotic extracts, which was abolished upon treatment of extracts with the Cdk1 inhibitor Purvalanol A. Taken together, these results show that importin α is phosphorylated at serine 62 during mitosis by Cdk1.

### Impaired importin α phosphorylation causes mitotic defects

We then sought to determine the effects of importin α phosphorylation on mitotic progression. To rule out potential chronic effects of expressing phospho-deficient mutant importin α, we generated stable normal rat kidney (NRK) cell lines that express human FLAG-tagged wild type importin α (hereafter referred to as “WT-expressing”) or mutant importin α S62A (hereafter referred to as “S62A-expressing”) upon doxycycline treatment. These importin α-expressing cells were synchronized and released from a thymidine G1/S block, and then their progression into M-phase was followed by phase contrast time-lapse microcopy. Average time to mitotic entry of WT- and S62A-expressing cells was comparable, indicating that ectopic expression of importin α does not impact interphase progression or the G2/M transition (Figure 2A). However, the time from cell rounding to cytokinesis onset (as a measure for the duration of mitosis) was prolonged by 13% (or 5 min) in S62A-expressing cells compared to WT-expressing cells (Figure 2B). Moreover, whereas WT-expressing cells divided normally, 21.2% of mutant cells with phospho-deficient importin α S62A exhibited cytokinesis defects, often generating more than two daughter cells (Figures 2C and 2D).

**Figure 2.**
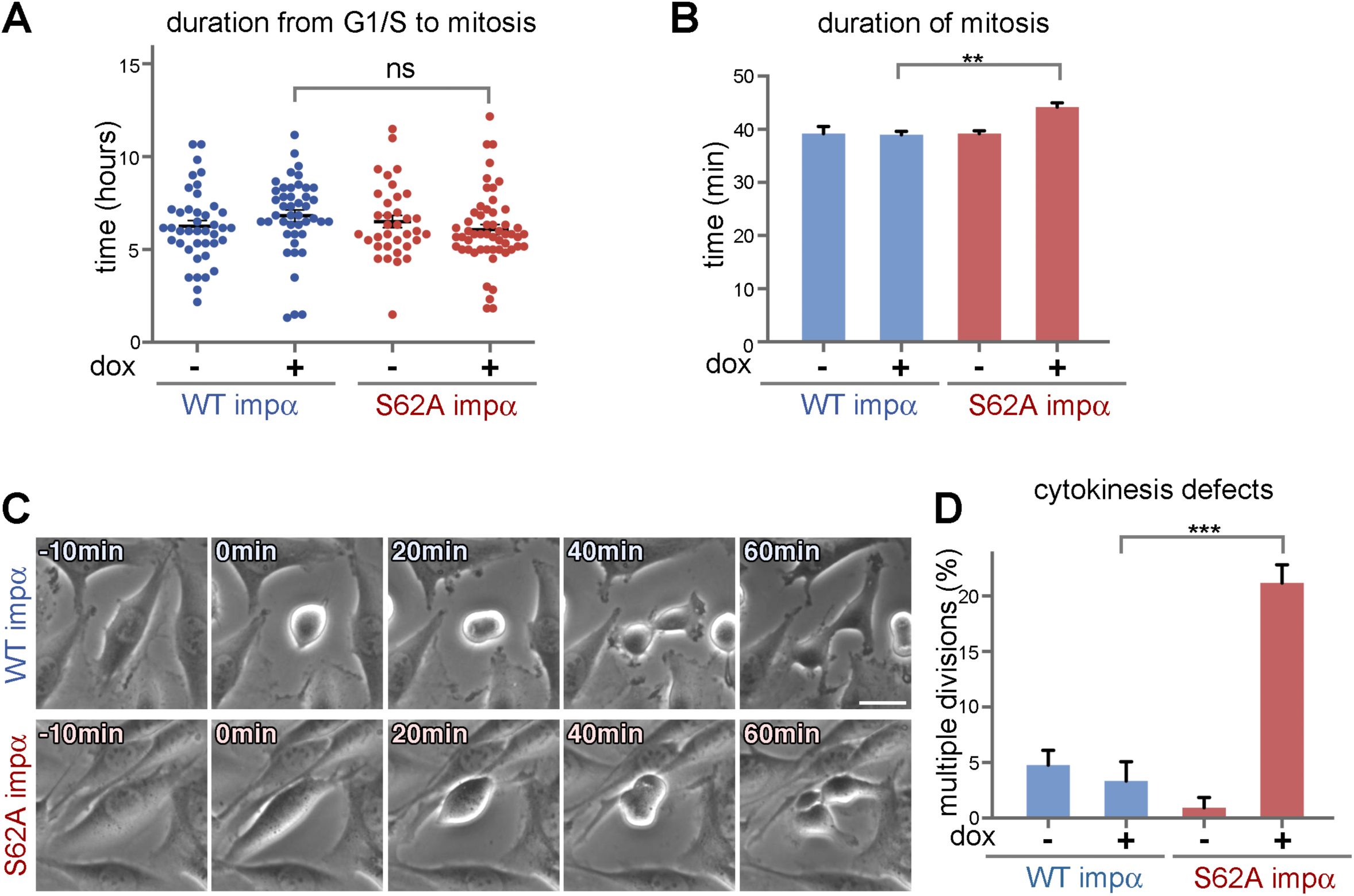
Importin α S62A induces mitotic defects. NRK cells non-induced or induced to express WT importin α or importin α S62A were synchronized at the G1/S transition by means of thymidine treatment for 16 h. The cells were then released from the thymidine block and cell cycle progression was monitored by time-lapse phase contrast microscopy. (A) Importin α expression does not affect mitotic entry. The duration between release from G1/S arrest to mitotic entry is plotted (n>90). Error bars represent SEM. (B) Importin α S62A-expressing cells exhibit prolonged mitotic duration (by 5 min; n>90). (C and D) NRK cells expressing importin α S62A exhibit cytokinesis defects (n>90). Error bars represent SEM. Scale bar = 20 µm.

### Mitotic phosphorylation of importin α controls spindle orientation

To further investigate the effects of mutant importin α S62A on mitosis, we analyzed the 3D-orientation of metaphase spindle using confocal microscopy. Cells arrested in metaphase were double-stained for α-tubulin and γ-tubulin to label spindle microtubules and mark the spindle poles, respectively. Then we measured spindle length and calculated spindle orientation along the z-axis (Figure 3A). We found that average distances between the two spindle poles did not differ significantly between WT- and S62A-expressing cells (Figures 3B and 3D), indicating that the overall structure of the spindle and the dynamics of spindle microtubule fibers were not affected (Baudoin and Cimini, 2018). Notably, in WT-expressing cells, both spindle poles were detected within the same z-section and were aligned parallel to the substratum. In contrast, the spindle poles of S62A-expressing cells were located in different z-planes, revealing that the spindle axis was tilted relative to the coverslip surface. The angle between the coverslip surface and the plane in which the two spindle poles resided showed an aberrant 13° tilt of the spindle axis in S62A-expressing cells (Figure 3C). These results indicate that expression of phospho-deficient importin α S62A alters the orientation of the mitotic spindle and affects cell division.

**Figure 3.**
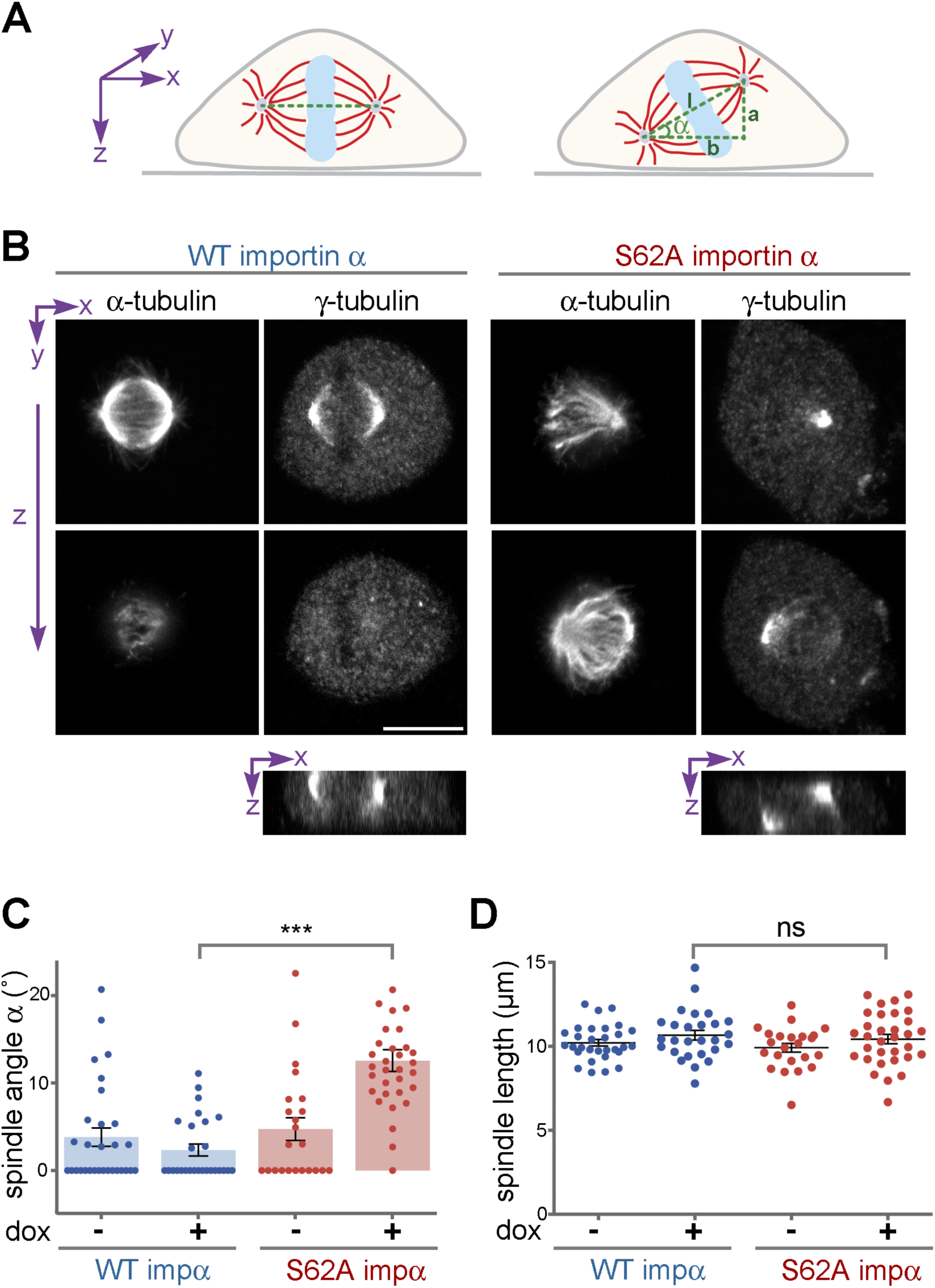
Phosphorylation of importin α controls spindle orientation. (A) Schematic illustration of spindle orientation in tissue culture cells. The axis of a normal spindle is oriented parallel to the coverslip, whereas the axis of a misoriented spindle is tilted. The angle α of spindle tilt was calculated as: α=arctan (b/a). Spindle length (l) was calculated as the 3D distance between the spindle poles (l^2^=a^2^+b^2^). (B) Importin α S62A causes misorientation of the spindle. NRK cells expressing WT importin α or importin α S62A were arrested at metaphase using MG132 and then dual-stained for α-tubulin and γ-tubulin to label spindle microtubules and the spindle poles, respectively. Z-sections were captured by confocal microscopy. Orthogonal sections through the z stacks along the x-axis (bottom panels) indicate the relative spindle pole localization on the z-axis. Scale bar = 10 μm. (C) Importin α S62A results in tilted spindles. The spindle tilt angle α was calculated as shown in (A). n>30, error bars represent SEM. (D) Importin α S62A does not alter spindle length. The spindle length l was determined as shown in (A). n>30, error bars represent SEM.

### Mutant importin α S62A inhibits astral microtubule growth

Proper orientation of the spindle depends on the astral microtubules that connect the spindle to the cell cortex (Lu and Johnston, 2013). To decipher if astral microtubules in cells expressing mutant importin α S62A were compromised, we arrested the cells in metaphase using MG132 and then investigated the microtubule networks by confocal microscopy. To better visualize the astral microtubules, which are less abundant and dimmer compared to interpolar and kinetochore microtubules, we increased image exposure. We found that the astral microtubule network originating at steady-state spindle poles of S62A-expressing cells was less dense and the filaments were shorter relative to WT-expressing cells (Figure 4A). Furthermore, the relative fluorescence intensity of α-tubulin staining in the astral area of S62A-expressing cells was diminished by 30% compared to WT cells (reduced from 9% to 6%, Figure 4B).

**Figure 4.**
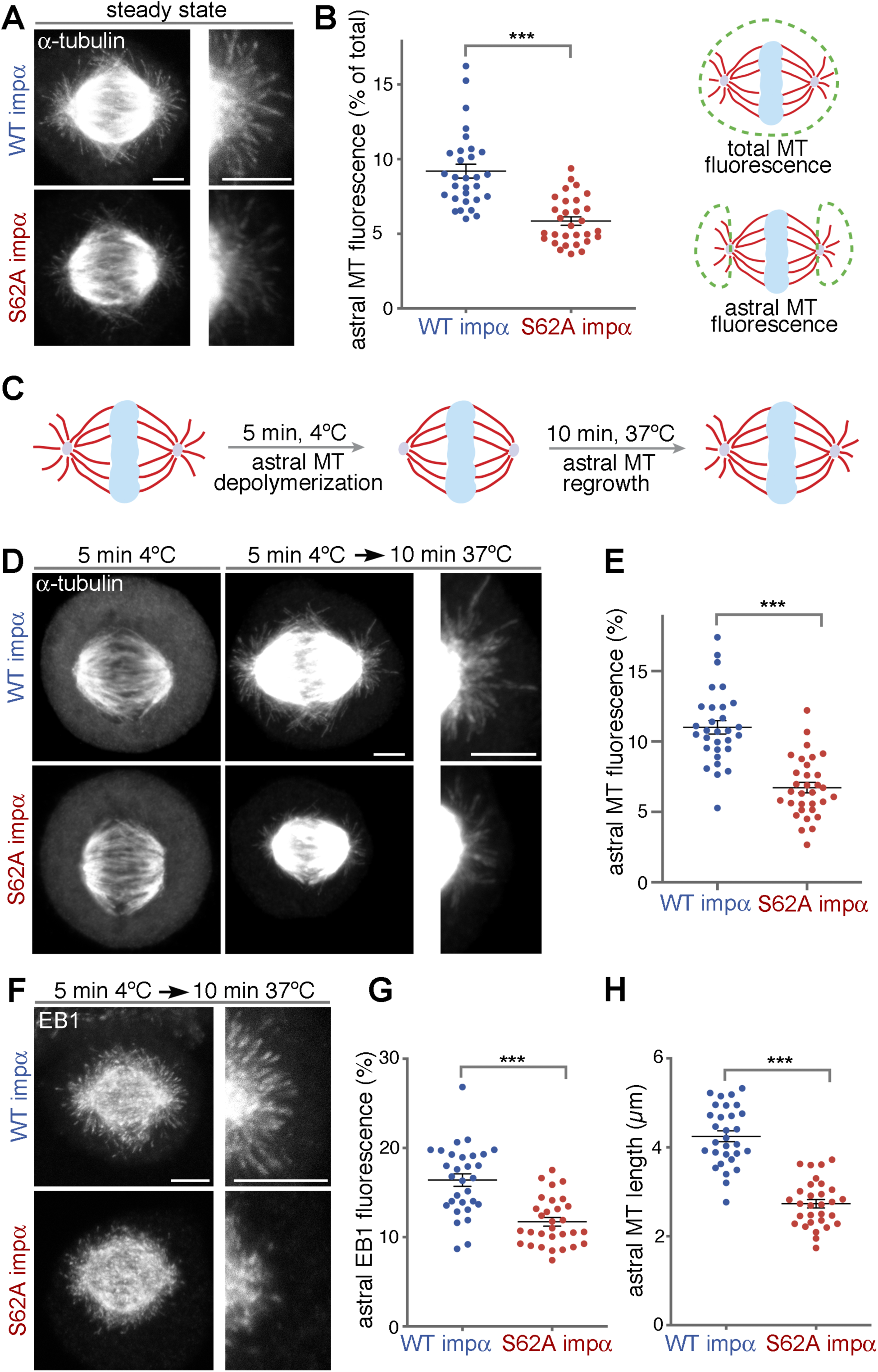
Importin α S62A inhibits microtubule growth. (A) Importin α S62A compromises astral microtubules. NRK cells expressing WT importin α or importin α S62A were arrested at metaphase using MG132 for 1 h and stained for α-tubulin. Maximum intensity projections of confocal z-sections are shown. Enlargements of the astral microtubule area are shown in the panels at right. (B) Quantitation of (A). Astral microtubule fluorescence was measured as the percentage of astral microtubule intensity relative to the total microtubule intensity, as shown in the schematic. n>30, error bars represent SEM. (C) Schematic illustration of the cold-induced astral microtubule regrowth assay. Astral microtubules were depolymerized by 5 min of cold treatment, and then allowed to regrow by shifting the cells back to 37 °C for 10 min. (D) Importin α S62A inhibits astral microtubule regrowth upon cold treatment. NRK cells arrested at metaphase using MG132 were placed on ice for 5 min to depolymerize astral microtubules but not spindle filaments. After 10 min regrowth at 37 °C, cells were fixed and stained for α tubulin. Confocal microscopy images represent maximum intensity projections. (E) Astral microtubule fluorescence was calculated as in (B). n>30, error bars represent SEM. (F) Importin α S62A inhibits EB1 localization to the astral area. EB1 was stained following the procedure in (C and D). Maximum intensity projections are shown. (G) EB1 fluorescence was calculated as for astral microtubule intensity. n>30, error bars represent SEM. (H) Importin α S62A shortens astral microtubule length. The lengths of astral microtubules were determined as the distance between the tip of an EB1 signal comet and the spindle pole. For each cell, the average length of the three longest filaments from each pole was used to represent nascent microtubule length. n>30, error bars represent SEM. All scale bars = 5 µm.

To determine if the S62A mutation affects initiation of astral microtubules, we performed a microtubule regrowth assay upon cold-induced depolymerization (Figure 4C). Cells arrested in metaphase were placed on ice for 5 min, resulting in full depolymerization of astral microtubules (Figure 4D). The cells were then shifted to 37 °C for 10 min to allow microtubule regrowth and stained for α-tubulin. This assay showed that the astral microtubule network in WT-expressing cells was restored to steady-state conditions upon temperature shift, whereas astral microtubule regrowth was reduced by 25% in S62A-expressing cells (Figure 4E).

To further test if the shorter and less abundant astral microtubule network in S62A-expressing cells was due to diminished microtubule growth, we performed another microtubule regrowth assay and stained for EB1 (Figure 4F). EB1 preferentially localizes at the tips of growing microtubules and functions as a marker for growing microtubule filaments. Consistent with the tubulin staining (Figure 4E), EB1 signal intensity on astral filaments in S62A-expressing cells was reduced by 29% (from 16.4% to 11.7% of the EB1 signal in the astral area) (Figure 4G). Furthermore, the EB1 comets did not emanate as far from the spindle poles of S62A-expressing cells (2.7 µm) as it did in WT-expressing cells (4.2 μm) (Figure 4H). These results demonstrate that expression of mutant importin α S62A impairs astral microtubule growth and dynamics, resulting in spindle misorientation.

### TPX2 is required for astral microtubule formation

During mitosis, importin α silences the activity of the spindle assembly factor TPX2 in the cytoplasm. Upon release from importin α inhibition, TPX2 binds to and activates Aurora A, an essential kinase for spindle assembly (Garrido and Vernos, 2016). Previously, we demonstrated that mitotic Golgi membranes can initiate localized microtubule assembly by liberating TPX2 from importin α inhibition (Wei et al., 2015). Given that mitotic Golgi membranes accumulate around the spindle poles (Wei and Seemann, 2009a), we wondered if TPX2 binding to Aurora A drives astral microtubule formation. To test this possibility, we took advantage of AurkinA, a recently developed small-molecule inhibitor that specifically blocks the interaction of TPX2 with Aurora A (Figure 5A) (Janeček et al., 2016). We first tested the overall efficacy of the inhibitor on the spindle microtubules. Cells arrested in metaphase were treated with 200 µM AurkinA for 0, 15 or 30 min, and then immuno-stained for tubulin (Figure 5B). We found that prolonged AurkinA treatment (for 30 min) weakened the microtubule filament network, whereas 15-min AurkinA incubation mainly impacted astral microtubules compared to spindle filaments (Figures 5C and 5D). Furthermore, the 15 min AurkinA treatment reduced the lengths of astral microtubules, as well as EB1 signal intensity and density in the astral microtubule area (Figures 5E and 5F). These results indicate that TPX2 binding to Aurora A kinase is required for astral microtubule formation.

**Figure 5.**
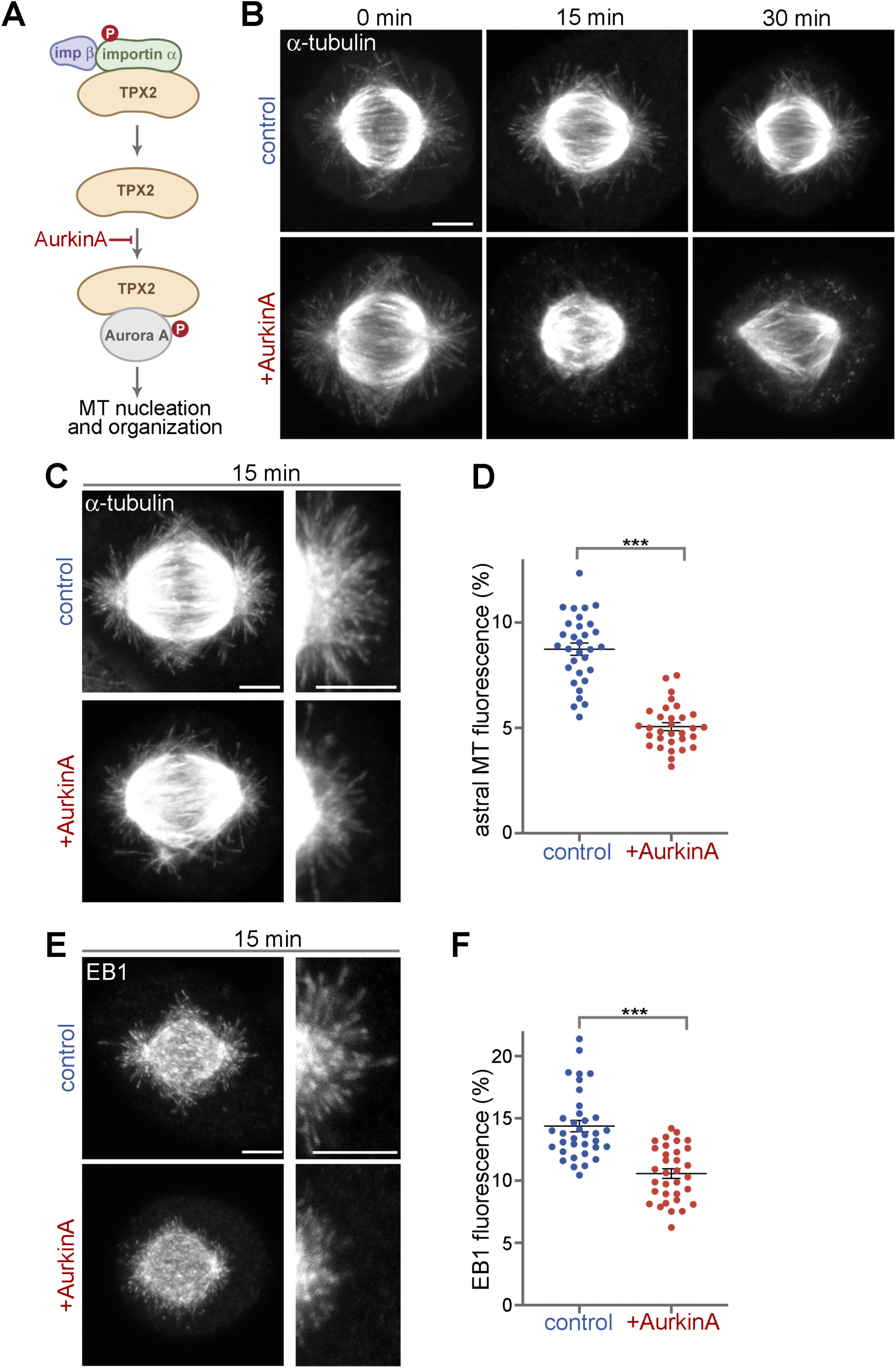
TPX2 is required for astral microtubule formation. (A) AurkinA is a small molecule inhibitor that specifically blocks the interaction of TPX2 with Aurora A kinase. (B) AurkinA alters spindle organization. NRK cells arrested at metaphase using MG132 for 1 h were treated with AurkinA for 0, 15 or 30 min and then fixed and stained for α-tubulin. Maximum intensity projections are shown. (C) AurkinA compromises astral microtubules. After 15 min of AurkinA treatment, NRK cells were fixed and stained for α-tubulin. (D) Quantitation of (C). n>30, error bars represent SEM. (E) AurkinA inhibits astral microtubule growth. After 15 min of Aurkin A treatment, NRK cells were fixed and stained for EB1. (F) Quantitation of (E). n>30, error bars represent SEM. All scale bars = 5 µm. Maximum intensity projections of confocal z-sections are shown.

### GM130 regulates astral microtubule growth via phosphorylated importin α

Upon entry into M-phase, the Golgi ribbon is swiftly disassembled into a collection of vesicles and clusters of membranes (termed mitotic Golgi clusters). By metaphase, these mitotic Golgi clusters are concentrated around the two spindle poles in the vicinity of astral microtubule filaments (Jokitalo et al., 2001). Competition for importin α binding by GM130 on Golgi membranes activates the microtubule initiation role of TPX2 (Wei et al., 2015). Therefore, we tested if the mitotic Golgi clusters can initiate astral microtubule polymerization through TPX2 and if, in turn, the astral microtubules are important for proper localization of mitotic Golgi clusters. We found that expression of WT or S62A-mutant importin α in interphase NRK cells did not affect the morphology or localization of the Golgi apparatus in the perinuclear region of those cells (Figure 6A). However, during mitosis, we observed that expression of importin α S62A not only diminished astral microtubules (as shown in Figure 4), but mitotic Golgi clusters were also displaced from the spindle pole area and were reduced in size in S62A-expressing cells (Figure 6B).

**Figure 6.**
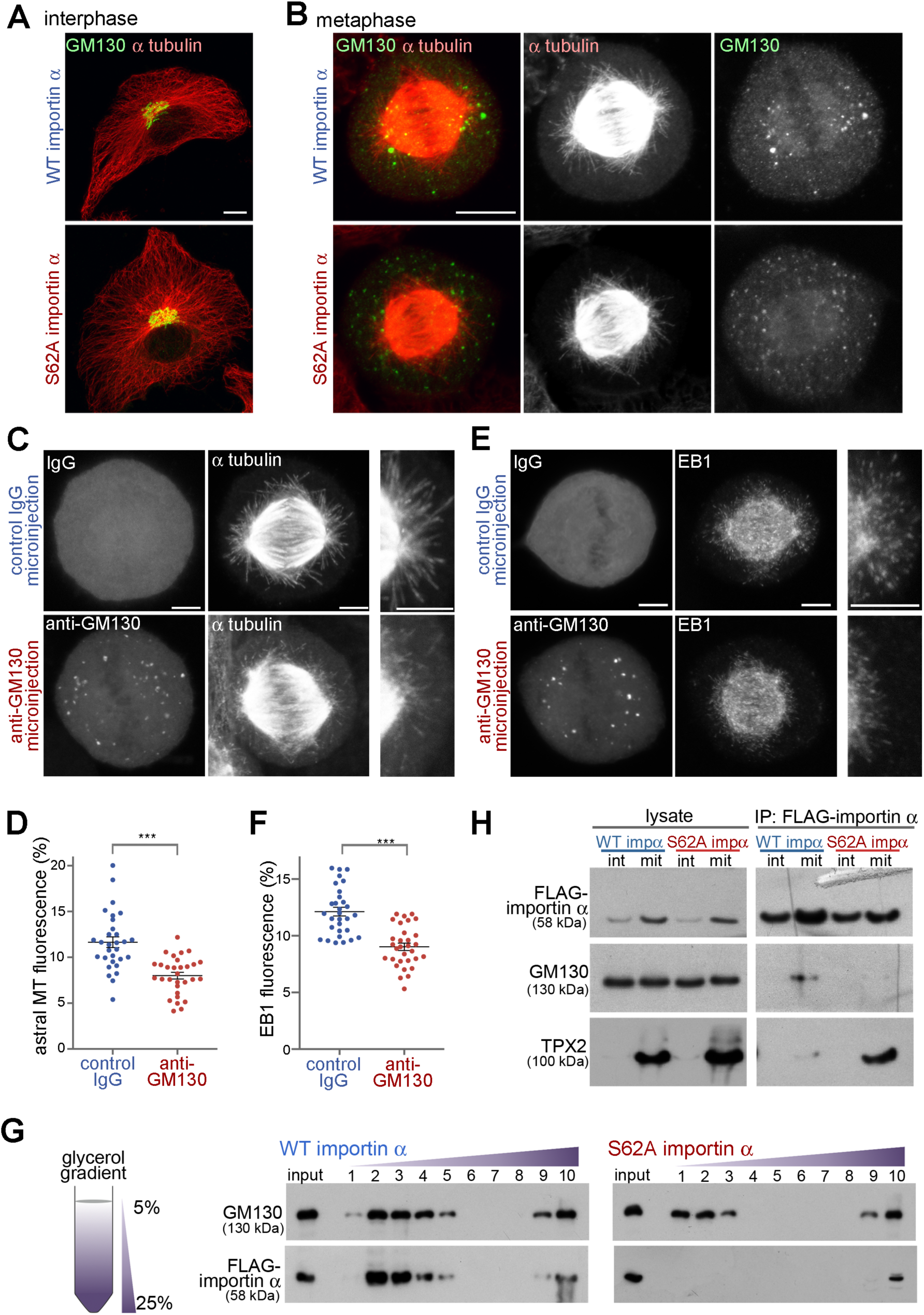
GM130 regulates astral microtubule growth via phosphorylated importin α. (A) Importin α S62A does not alter Golgi structure or the microtubule network during interphase. Interphase NRK cells expressing WT importin α or importin α S62A were fixed and stained for α-tubulin and the Golgi membrane protein GM130. Maximum intensity projections are shown. Scale bar = 10 µm. (B) Importin α S62A delocalizes mitotic Golgi clusters from the spindle pole area. NRK cells expressing WT importin α or importin α S62A were arrested at metaphase using MG132, and then fixed and stained for GM130 and α-tubulin. Maximum intensity projections are shown. Scale bar =10 µm. (C and E) Blocking the GM130-importin α interaction inhibits astral microtubule growth and delocalizes Golgi clusters from the spindle poles. NRK cells were arrested at metaphase for 1 h and microinjected with inhibitory antibody against GM130 or with control IgG. Injected cells were incubated for another 30 min, followed by cold treatment for 5 min. They were then allowed to regrow microtubules for 10 min, before fixation and staining for α-tubulin, EB1 and injected IgG. Maximum intensity projections are shown. Scale bars = 5 µm. (D and F) Quantitation of astral microtubule fluorescence and astral EB1 fluorescence from (C and E). n>30, error bars represent SEM. (G) Importin α S62A does not associate with mitotic Golgi membranes. Post-chromosomal supernatant of mitotic HeLa cells expressing WT importin α or importin α S62A was centrifuged through a linear 5%-25% glycerol gradient to separate membrane components by size. Ten fractions were collected from the top. The membranes from each fraction were collected by centrifugation and then analyzed by Western blotting. (H) Phosphorylation of importin α shifts its binding preference from TPX2 to GM130. Interphase or mitotic HeLa cells expressing FLAG-tagged WT importin α or importin α S62A were lysed and incubated with anti-FLAG antibody-conjugated beads. Beads were then washed with lysis buffer and analyzed by Western blotting.

We then sought to acutely interfere with the microtubule initiation capacity of the mitotic Golgi clusters. The microtubule initiation activity of GM130 can be blocked by microinjection of inhibitory anti-GM130 antibodies (Wei et al., 2015). We microinjected affinity-purified GM130 antibodies or control IgG into MG132-arrested metaphase NRK cells. Then, 30 min after the injection, we depolymerized astral microtubules by cold treatment, before shifting the cells to 37 °C for 10 min to allow filament regrowth (Figure 6C). The cells were then stained for the injected antibodies (to discriminate the cells), as well as for tubulin and EB1. Blocking GM130 activity phenocopied the effects of AurkinA treatment (Figure 5) by attenuating astral microtubules (Figures 6C and 6D) and their growth dynamics (as revealed by reduced EB1 labeling; Figures 6E and 6F). Consistent with expression of mutant importin α S62A, we further noted from the staining patterns of injected GM130 antibodies that Golgi clusters were more fragmented and delocalized from the spindle pole area (Figures 6C and 6E). These results suggest that the GM130-TPX2 pathway activates astral microtubule assembly and is enabled by importin α phosphorylation.

Importin α is recruited to Golgi membranes during mitosis via GM130 (Wei et al., 2015). If mitotic phosphorylation of importin α is required for binding to GM130, then importin α S62A should not associate with mitotic Golgi membranes. We tested this possibility using velocity gradient centrifugation to separate smaller mitotic Golgi vesicles from larger membranes (Jesch et al., 2001; Seemann et al., 2002). HeLa cells arrested in mitosis by means of the Eg5 kinesin inhibitor S-Trityl-L-Cysteine were collected by shake-off and mechanically lysed using a ball bearing homogenizer. The post-chromosomal supernatant was then centrifuged through a linear 5%-25% glycerol gradient and then the membranes of collected fractions were immunoblotted for GM130 and FLAG-tagged importin α (Figure 6G). Consistent with our previous report (Wei et al., 2015), WT importin α co-migrated with GM130 in the top fractions of the gradient, demonstrating that importin α had been recruited to mitotic Golgi membranes. In contrast, phospho-deficient importin α S62A did not associate with mitotic Golgi membranes in the top fractions of the gradient. We also observed that the GM130-containing membrane fractions were shifted upwards to the lighter membranes in the gradient, in line with our immunofluorescence data showing that smaller mitotic Golgi clusters formed in S62A-expressing cells (Figure 6B). Some of the importin α remained in the bottom fraction of the gradient comprising the bulk of the cellular membranes, consistent with previous reports that importin α also associates with the nuclear envelope and the plasma membrane during mitosis (Hachet et al., 2004; Brownlee and Heald, 2019).

To further assess the phosphorylation dependency of the importin α-substrate interaction, we pulled down FLAG-tagged WT or importin α S62A from mitotic lysates and probed for association of endogenous GM130 and TPX2 by means of immunoblotting (Figure 6H). The results showed that TPX2 binds much more strongly to importin α S62A than to phosphorylated WT importin α. Conversely, binding of GM130 to phosphorylated WT importin α was greatly enhanced relative to phospho-deficient importin α S62A. These results evidence that phosphorylation of importin α shifts its substrate preference from TPX2 to GM130, allowing GM130 on mitotic Golgi membranes to sequester importin α from TPX2 and activate Aurora A to regulate astral microtubules and spindle orientation.

## Discussion

Here, we report that TPX2 activation regulates astral microtubule dynamics and is essential for correct spindle orientation (Figure 7). Upon mitotic entry, phosphorylation of importin α at Ser62 by Cdk1 induces a switch in substrate preference from TPX2 to GM130 and thereby enables GM130 to activate TPX2 through direct competition. The phospho-incompetent importin α S62A point mutant prevents GM130 from activating TPX2, reducing astral microtubules and causing spindle misorientation. Furthermore, the characteristic clustering of mitotic Golgi membranes at the spindle poles is astral microtubule-dependent and requires TPX2 activation.

**Figure 7.**
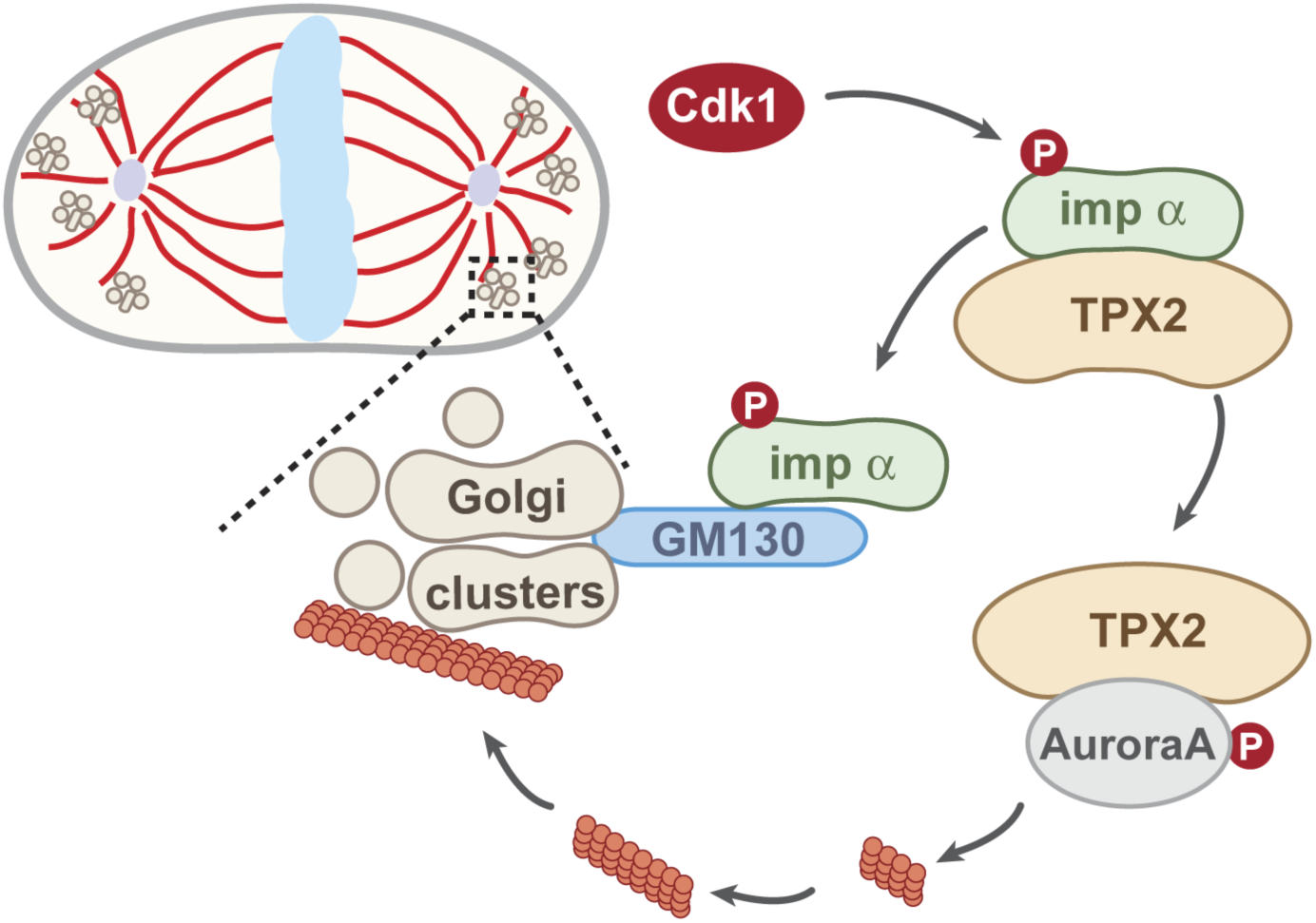
Model of the TPX2-mediated astral microtubule assembly pathway. Phosphorylation of importin α at Ser62 by Cdk1 switches its substrate preference from TPX2 to GM130. This switch enables GM130 on the mitotic Golgi clusters at the spindle poles to locally sequester importin α from TPX2 through direct competition. TPX2 is thereby relieved from inhibition, allowing it to bind to and activate Aurora A kinase to drive astral microtubule assembly and facilitate spindle orientation.

Spindle orientation is important in determining cell fate and function. For example, in mammalian skin epidermal cells, the orientation of the spindle defines the division plane and, consequently, cell fate (Barr and Gruneberg, 2017). Embryonic epidermal progenitor cells initially proliferate in a single layer by dividing parallel to the basement membrane. During stratification, the cells may also divide perpendicular to the basement membrane, generating one daughter cell that remains proliferative in the basal layer and another that is destined for terminal differentiation. Therefore, spindle orientation must be tightly controlled, as defects in spindle orientation trigger unbalanced growth and differentiation, outcomes linked to hyperproliferation and cancer (Ragkousi and Gibson, 2014; Asare et al., 2017). Similarly, aberrant spindle orientation due to impairment of astral microtubule-cortex connectivity in *Drosophila* alters the balance between neural progenitor expansion versus differentiation, leading to defects in neuroblast homeostasis (Cabernard and Doe, 2009). Moreover, knockdown of ASPM (abnormal spindle-like microcephaly-associated gene), which directly contributes to the growth of astral microtubules, causes abnormal spindle orientation in neuroepithelial progenitor cells, and mutation of ASPM is the most common defect observed in human autosomal recessive primary microcephaly (Fish et al., 2006; Gai et al., 2016). Interestingly, similar defects have been reported for GM130 mutations *in vivo*. Knockdown of GM130 in Zebrafish leads to microcephaly (Shamseldin et al., 2016). The Purkinje cells of GM130-deficient mice exhibit neurodegeneration (Liu et al., 2016), and a patient with a homozygous loss-of-function mutation of GM130 developed neuromuscular disorder including progressive microcephaly (Shamseldin et al., 2016). These phenotypes have been broadly assigned to defective secretory pathways in which GM130 functions as a vesicle tethering factor during interphase. However, our present work shows that GM130 regulates spindle orientation, suggesting that deficiencies in GM130 mitotic function might further contribute to these neuronal disorders through impaired cell division and development.

Ran-dependent TPX2 activation takes place at the chromosomes. Analysis of RanGTP distribution in mitotic cells detected a low concentration of RanGTP in the astral area where Ran-mediated activation of TPX2 is unlikely to happen (Kaláb et al., 2006). Therefore, initiation and growth of astral microtubules has often been attributed solely to the centrosomes and microtubule-stabilizing proteins. Our findings have identified TPX2 as a key molecule in promoting astral microtubule growth. The direct competition-based activation of TPX2 on Golgi clusters does not require RanGTP and facilitates microtubule organization in a distinct spatiotemporal fashion.

Our results also indicate that phosphorylation-deficient importin α S62A inhibits TPX2 from being activated by GM130 at the spindle poles, leading to diminished astral microtubule density that, in turn, misorients the spindle. Since importin α modulates several mitotic regulators, other mitotic events could also be affected by importin α S62A, such as the cytokinesis defects we report here. However, our observed phenotype of spindle misorientation induced by importin α S62A should mainly arise from compromised astral microtubule dynamics. We did not record spindle oscillation, which has often been reported for cells with spindle-cell cortex junction defects (Kotak and Gönczy, 2014; Petry, 2016). The centrosomal protein ASPM targets the citron kinase CITK to the spindle poles, where it specifically regulates astral microtubule dynamics (Gai et al., 2016). In ASPM-knockdown cells, spindle length is unchanged and spindle oscillation has not been observed. Moreover, the metaphase to anaphase transition is delayed by five minutes in such cells and the spindle is misoriented (Gai et al., 2016). Expression of importin α S62A essentially phenocopied knockdown of ASPM. Therefore, it is reasonable to conclude that importin α regulates astral microtubule growth by promoting astral microtubule dynamics from the Golgi clusters at the pole area. Moreover, our acute inhibition of the GM130-TPX2 pathway slowed down microtubule growth, prompting us to conclude that the GM130-TPX2 pathway is an important element of the mechanism controlling astral microtubule dynamics.

Apart from the Golgi, other membrane components have been reported to influence astral microtubule dynamics (Barr and Gruneberg, 2017). Recycling endosomes can promote microtubule growth in mitotic extracts, and knockdown of the recycling endosomal protein Rab11 results in compromised astral microtubules and rotating spindles (Hehnly and Doxsey, 2014). Based on those findings, it has been proposed that the poleward movement of recycling endosomes transports microtubule-nucleating material like γ-tubulin and GCP4 protein to the centrosomes and facilitates the polymerization of astral microtubules. Peroxisomes also play a role in orienting the spindle. Knockdown of peroxisomal protein Pex11b results in diminished cell cortex localization of NuMA, contributing to spindle oscillation and misorientation (Asare et al., 2017). Here, we report mitotic Golgi clusters as being membrane organelles that regulate spindle orientation. We showed in a previous study that the association between mitotic Golgi membranes and the spindle is required for a single Golgi ribbon to reform upon mitotic exit (Wei and Seemann, 2009a). In the current study, we have demonstrated that the association of mitotic Golgi clusters with astral microtubules originates from the ability of the former to nucleate microtubules. When we blocked GM130-triggered TPX2 activation by microinjecting inhibitory antibodies, the Golgi clusters became delocalized from the spindle poles and dispersed. In addition, expression of importin α S62A diminishes the size of these Golgi clusters. Although it is still not clear how mitotic Golgi clusters are organized *in vivo*, our results suggest that the microtubule network plays an integral role in maintaining the morphology and localization of the mitotic Golgi clusters at the spindle poles.

Post-translational modifications of importin family proteins affect their binding affinity to their substrates, influencing the speed of cargo transport into the nucleus during interphase. Importin α is repurposed as a spindle modulator during mitosis, but it was not clear how it specifically recognizes spindle assembly factors at this time given the amount of NLS-containing proteins that are released into the cytoplasm upon nuclear envelope breakdown (Giesecke and Stewart, 2010). Our study shows that Cdk1-mediated and mitotic-specific phosphorylation of importin α at Ser62 shifts its binding preference from TPX2 to GM130. Since importin α regulates other proteins that play important roles in mitosis, phosphorylation of importin α potentially also modulates their activities. While we were preparing this manuscript, it was reported that Ser62 phosphorylation is correlated with decreased binding to TPX2, but the mechanistic significance at the cellular level was not explored (Guo et al., 2019). A recent report based on the structure of non-phosphorylated importin α complexed with the N-terminal fragment of GM130 suggests that, unlike other NLS sequences, GM130 competes with TPX2 for the minor binding site of the NLS-binding pockets in importin α, thereby preventing other NLS-containing cargos from activating TPX2 by competitive inhibition (Chang et al., 2019). A more thorough study on phosphorylation-induced conformational change should help further establish the exact mechanism by which importin α is regulated and repurposed throughout the cell cycle.

## Materials and methods

### Plasmids and cell lines

Human importin α1 (KPNA2) WT or the S62A point mutant was cloned into pET30a for His-importin α recombinant protein purification. The S62A mutation was generated by quick-change PCR mutagenesis. Importin α with an N-terminal FLAG tag was cloned into pLVX-TetOne-Puro (Clontech) to generate stable cell lines. pLVX-FLAG-KPNA2 WT or S62A was co-transfected with psPAX and pVSVG into HEK293T cells to produce lenti-virus, which was then used to transduce NRK and HeLa cells. Two days later, we added 5 μg/ml puromycin (RPI) and stable clones were selected for seven days. Individual colonies were picked and single clones were isolated by limited dilution. Expression of FLAG-importin α WT or S62A was induced for 16 h by addition of 1 µg/ml doxycycline (Sigma).

### Cell culture and drug treatments

HEK293T, HeLa and normal rat kidney (NRK) cells were cultured at 37 °C and 5% CO_2_ in complete growth medium [Dulbecco’s modified Eagle’s medium (Mediatech) supplemented with 10% cosmic calf serum (HyClone), 100 units/ml penicillin and 100 µg/ml streptomycin]. NRK cells were synchronized at the G1/S phase transition by 16 h treatment with 2 mM thymidine (Chem-Impex). Then, we washed out the thymidine and added 24 µM 2’-deoxycytidine (Chem-Impex). Five hours after release from thymidine, we added 10 µM MG132 (Boston Biotech) for 1 h to arrest cells in metaphase.

HeLa cells and HeLa cells expressing FLAG-tagged WT or S62A importin α were synchronized in mitosis by treatment with 20 µM S-Trityl-L-Cysteine (Acros) for 16 h. The mitotic cells were then collected by shake-off, followed by centrifugation. We employed 200 µM AurkinA (Aobious) in complete culture medium to block the association of TPX2 with Aurora A kinase.

### Antibodies

For Western blotting, we used mouse monoclonal antibodies against importin α (BD Transduction Lab, Cat. #610485, 25 ng/ml) and GAPDH (GA1R, Invitrogen, Cat. #MA5-15738, 20 ng/ml), as well as rabbit polyclonal antibodies against Phospho-Histone H3 (Ser10) (Millipore, Cat. #06-570, 25 ng/ml), TPX2 (Proteintech, Cat. #11741-1-AP, 150 ng/ml), FLAG-tag (Sigma, Cat. #F7425, 100 ng/ml), and GRASP55 (Rich, serum 1:5000) (Wei and Seemann, 2009a).

For immunofluorescence, we used mouse monoclonal antibodies against EB1 (BD Transduction Lab, Cat. #61534, 1 μg/ml), α-tubulin (TAT1; supernatant 1:200) (Woods et al., 1989), and GM130 (NN2C10, 5 µg/ml, used in Figures 6A and 6B) (Seemann et al., 2000), rabbit polyclonal antibody against γ-tubulin (Sigma, Cat.# T3559), as well as rat monoclonal antibody against α-tubulin (Chemicon, Cat. #MAB1860, 5 µg/ml). For microinjection, we used affinity-purified rabbit anti-GM130 (Wei et al., 2015) and rabbit IgG (Sigma).

Secondary antibodies were as follows: Alexa Fluor 488- or Alexa Fluor 594-conjugated to goat anti-mouse or goat anti-rabbit IgG (Invitrogen), Alexa Fluor 568-conjugated to goat anti-rat IgG (Invitrogen), HRP-conjugated goat anti-rabbit (Jackson ImmunoResearch), HRP-conjugated goat anti-mouse IgG (Invitrogen).

### Microinjection

NRK cells grown on glass coverslips were microinjected using a FemtoJet system (Eppendorf) and a micromanipulator 5171 (Eppendorf) attached to an Eclipse TE300 inverted microscope (Nikon). The cells were switched to complete growth medium supplemented with 50 mM HEPES/KOH pH 7.4 before injections. The cells were microinjected with 2 mg/ml affinity-purified rabbit anti-GM130 or 2 mg/ml control IgG in H/K buffer (20 mM HEPES/KOH pH 7.4, 50 mM KOAc). Injected cells were incubated at 37 °C for 30 min and then subjected to an astral regrowth assay. Cells were then fixed in −20 °C methanol for EB1 staining or in microtubule-stabilizing fixative to stain for α-tubulin.

### Immunofluorescence

For the astral microtubule regrowth assay, we replaced the medium with cold complete growth medium for 5 min to depolymerize astral microtubules and then changed it back to 37 °C complete growth medium and continued incubating the cells at 37 °C for 10 min to allow microtubule regrowth. To stain for aster microtubules, the cells were fixed for 20 min at 37 °C in microtubule-stabilizing fixative (3.7% formaldehyde, 0.1% glutaraldehyde, 60 mM PIPES, 25 mM HEPES, 2 mM MgCl_2_ and 10 mM EGTA, pH 6.9). The reaction was quenched for 5 min with sodium borohydride in 20 mM Tris pH 7.4 and 150 mM NaCl, before being permeabilized, washed three times with TBS-Tx (20 mM Tris pH 7.4, 150 mM NaCl and 0.1% Triton X-100), and blocked with 1 mg/ml BSA in TBS-Tx at room temperature for 30 min.

For all other immunofluorescence stainings, the cells were fixed and permeabilized for 15 min in methanol at −20 °C, then incubated with respective antibodies at 37 °C for 30 min, followed by Alexa-Fluor-conjugated secondary antibodies. DNA was stained for 10 min at room temperature with 1 µg/ml Hoechst 33342 (Invitrogen) in PBS and cells were embedded in Mowiol (Calbiochem) mounting solution (Wei and Seemann, 2009b).

### Subcellular fractionation

Mitotic HeLa cells expressing WT or importin α S62A were mechanically lysed using a ball bearing homogenizer with a clearance of 12 µm (Isobiotec). The post-chromosomal supernatant (PCS) was collected by centrifugation for 5 min at 1000 *g* at 4 °C, loaded onto a continuous 5%-25% glycerol gradient, and centrifuged for 30 min in a SW55Ti rotor (Beckman) at 35,000 rpm at 4 °C. Ten 0.5 ml fractions were collected from the top of the gradient, diluted with 1 ml of 20 mM HEPES/KOH pH 7.4, and membranes were collected by centrifugation for 30 min at 100,000 *g* at 4 °C. The supernatant was discarded, and the membrane pellets were resuspended in SDS sample buffer for further analysis by Western blotting.

### Co-immunoprecipitation

Interphase or mitotic HeLa cells expressing FLAG-importin α WT or S62A mutants were lysed for 20 min at 4 °C in lysis buffer [10 mM Tris pH 7.4, 150 mM NaCl, 1 mM EDTA, 1% TritonX-100, cOmplete protease inhibitor cocktail (Roche), 1 mM DTT, 10 mM β-glycerophosphate, 1 mM Na_3_VO_4_ and 10 mM NaF]. Lysates were cleared by centrifugation at 16,000 *g* for 10 min at 4 °C and then incubated with 10 µl anti-FLAG M2-Sepharose beads (Sigma) for 60 min at 4 °C. Beads were then washed three times with lysis buffer and boiled in SDS sample buffer to elute proteins for Western blotting analysis. For mass spectrometry, FLAG-importin α purified from mitotic and interphase HeLa cells was eluted from the beads using 300 µg/ml 3xFLAG-tag peptide (Apexbio). The 58 kDa bands were cut from the gel and processed by the Proteomics Core Facility at UT Southwestern.

### Protein purification

BL21(DE3) competent *E. coli* were transformed with pET30a-importin α (WT, S62A). Single clones were inoculated into 20 ml LB medium and grown overnight at 37 °C. The 20 ml cultures were then inoculated into 1000 ml LB to allow the bacteria to grow to OD_600_=0.6-0.8. Protein expression was induced by addition of 0.2 mM IPTG and incubation for 18 h at 16 °C. Bacteria were collected by centrifugation, resuspended in high salt buffer (HSB) [PBS supplemented with 1 M NaCl, cOmplete protease inhibitor cocktail (Roche), 1 mM DTT, 1 mM PMSF] and lysed using a French Press. Nickel-NTA agarose beads (Qiagen) were used to purify His-importin α (1 ml slurry for 1000 ml culture). The beads were then washed with HSB and directly used for *in vitro* phosphorylation (5 μl slurry per experiment).

### *In-vitro* phosphorylation assay

Mitotic HeLa cells were lysed for 20 min at 4 °C in lysis buffer [10 mM Tris pH 7.4, 150 mM NaCl, 1 mM EDTA, 1% TritonX-100, cOmplete protease inhibitor cocktail (Roche), 1 mM DTT, 10 mM β-glycerophosphate, 1 mM Na_3_VO_4_ and 10 mM NaF], and then the lysates were cleared by centrifugation at 16,000 *g* for 10 min at 4 °C. Nickel beads coated with recombinant His-tagged importin α were incubated at 30 °C for 30 min with pre-cleared mitotic lysate supplemented with 2 mM ATP pH 7.2. The Cdk1 kinase inhibitor Purvalanol A (LC Labs) was used at a concentration of 0.5 mM. Beads were then washed three times with HBS, eluted by boiling in SDS sample buffer, and analyzed by Phos-tag biotin blotting.

### Phos-tag gel and Phos-tag biotin

To detect importin α phosphorylation via *in vitro* phosphorylation assay, protein samples were separated by SDS-PAGE on a 10% gel and transferred to PVDF membrane. The PVDF membrane was then washed with TBST for 20 min before incubation with Phos-tag biotin solution [TBST supplemented with 6 µM ZnCl_2_, 2 µM Phos-tag Biotin (ApexBio) and 2 ng/ml streptavidin-HRP (Jackson ImmunoResearch)]. After 30 min incubation, the blot was developed by ECL detection.

To detect importin α phosphorylation on Phos-tag gels, mitotic or interphase cells were lysed for 20 min at 4 °C in lysis buffer (without EDTA) and cleared by centrifugation at 16,000 *g* for 10 min at 4 °C. For phosphatase treatment, 40 µl of the cleared lysates were incubated with 1 µl Lambda Protein Phosphatase (λ phosphatase) (New England Biolabs), 5 µl 10x buffer for Protein MetalloPhosphatases (New England Biolabs), and 5 µl 10 mM MnCl_2_ for 30 min at 30 °C. Proteins were separated on a 10% SDS gel containing 20 μM Phos-tag acrylamide (ApexBio) and 100 µM MnCl_2_. The gel was then washed for 1 h in transfer buffer containing 1 mM EDTA to remove the manganese. Proteins were subsequently transferred to PVDF membrane for immunoblotting.

### Microscopy and quantitation

Confocal microscopy was performed using a Zeiss LSM780 inverted microscope in combination with a Plan-Apochromat 63x/1.4 objective. Z-sections were captured at 0.5 µm intervals and maximum intensity projections are presented. Spindle angle was calculated using the formula α = arctan (b/a), where b = 0.5 μm multiplied by the number of z sections between the spindle poles and a = the distance between the projections of the two spindle poles on the x-y plane. Spindle length (l) was calculated as l^2^=a^2^+b^2^.

Relative fluorescence of astral microtubules was measured in ImageJ (as depicted in Figure 4B) as the fluorescence signal of the astral microtubule area/total microtubule fluorescence signal of the cell.

Astral microtubule length was determined by averaging the distance between the tip of the EB1 “comet” and the pole for the three longest filaments from each of the two asters.

Phase-contrast time-lapse microscopy was performed using an Axiovert 200M microscope (Zeiss) with an LD-A-PLAN 20x/0.3 Ph1 objective (Zeiss) and a Retiga 2000R CCD camera (QImaging) using MetaMorph 7.1.3 software (Molecular Devices). NRK cells expressing WT importin α or mutant importin α S62A released from thymidine blocking were changed to CO_2_-independent medium (Invitrogen), 10% cosmic calf serum (HyClone), 2 mM GultaMax (Invitrogen) and 100 units/ml penicillin and 100 μg/ml streptomycin. Phase contrast images were captured at 10 min intervals for 12 h, and cell cycle progression was analyzed using MetaMorph 7.1.3 (Molecular Devices) and ImageJ. The duration from G1/S phase to mitosis was determined as being from 30 min after thymidine washout until rounding of the cells. The duration of mitosis was measured as the time from rounding of the cells to onset of cytokinesis.

Image analysis was performed using ImageJ 2.0. Statistical analyses were conducted using Prism 8.3 software (GraphPad). Data shown are from three or more independent experiments. Error bars represent SEM. Statistical significance was assessed by Student’s t tests.

## Acknowledgements

We thank Yijun Zhang for comments on the manuscript and other suggestions, Gianni Guizzunti for insightful discussions, the Proteomics Core Facility for mass spectrometry analysis and the Light Microscopy Facility at the University of Texas Southwestern Medical Center for imaging support. J-H.W. is supported by an Academia Sinica Career Development Award (AS-CDA-109-L02). This work was supported by grants from the NIH (GM096070) and the Welch Foundation (I-1910) to J.S.

## Author contributions

H.G., J-H.W. and J.S. designed the project and wrote the manuscript. H.G. and J.S. performed the experiments, analyzed the data, and prepared the figures.

